# Multi-excitation Raman Spectroscopy Complements Whole Genome Sequencing for Rapid Detection of Bacterial Infection and Resistance in WHO Priority Pathogens

**DOI:** 10.1101/2022.02.08.479540

**Authors:** Adam Lister, Ekaterina Avershina, Jawad Ali, George Devitt, Niall Hanrahan, Callum Highmore, Jeremy Webb, Fredrik Muller, Sumeet Mahajan, Rafi Ahmad

## Abstract

Current methods for diagnosing acute and complex infections mostly rely on culture-based methods and, for biofilms, fluorescence *in-situ* hybridization. These techniques are labor-intensive and can take 2-4 days to return a test result, especially considering an extra culturing step required for the antibiotic susceptibility testing (AST). This places a significant burden on healthcare providers, delaying treatment and leading to adverse patient outcomes. Here, we report the complementary use of our newly developed multi-excitation Raman spectroscopy (ME-RS) method with whole-genome sequencing (WGS). Four WHO priority pathogens are AST phenotyped and their antimicrobial resistance (AMR) profile determined by WGS. On application of ME-RS method we find high correlation with the WGS characterization. Highly accurate classification based on the species (98.93%), wild-type/non-wild type (99.45%), and presence or absence of thick peptidoglycan layers in cell walls (100%), as well as at the individual strain level (99.29%). These results clearly demonstrate the potential of ME-RS as a rapid and first-stage tool for species, resistance and strain-level classification which can be followed up by WGS for confirmation. Such a workflow can facilitate efficient antimicrobial stewardship to handle and prevent the spread of AMR.

## Introduction

It has been more than 30 years since a new class of antibiotics was introduced to the market. This, together with the increased and inappropriate use of existing antibiotics means that we are heading towards a world in which many antibiotics are no longer effective. In addition to becoming the third leading cause of death^1^, antimicrobial resistance (AMR) has an enormous impact on worldwide economy. Each year the USA is losing US $55 billion and EU/EEA €1.6 billion due to the AMR^2^. In 2017 the World Health Organization (WHO) issued a report where the most critical pathogens were stratified into groups based on their threat for the increasing AMR spread and on urgency of action required^3^. ESKAPE pathogens (vancomycin-resistant *<u>E</u>nterococcus faecium* (VRE), methicillin-resistant and vancomycin-resistant *<u>S</u>taphylococcus aureus* (MRSA/VRSA), carbapenem-resistant and third-generation cephalosporin-resistant *<u>K</u>lebsiella pneumoniae, <u>A</u>cinetobacter baumannii, <u>P</u>seudomonas aeruginosa* and *<u>E</u>nterobacter* spp.) are included into the critical and high-priority pathogen groups. These pathogens have acquired resistance towards many antibiotics, including last-resort antibiotics such as carbapenems and colistin and are thus associated with high morbidity and mortality rates^4^. In 2019, 1.27 million people died due to resistant bacteria infections and 73% of them were caused by *E. coli, S. aureus, K. pneumoniae, S. pneumoniae, A. baumannii* and *P. aeruginosa*^1^.

A critical unmet need in the prevention of antimicrobial resistance (AMR) is rapid, accurate, and point-of-care (PoC) diagnosis to avoid the incorrect use of antibiotics. Without new diagnostics, the appropriate use of antibiotics and the treatment of patients with resistant bacterial infections will become increasingly challenging, compromising medical interventions such as surgeries, transplants, and chemotherapy. A recent retrospective study from a children’s hospital in Madrid (Spain) revealed that on giving at least one active antibiotic at the onset of bacteremia caused by carbapenem-resistant bacteria, the survival rate was increased by 80 % compared to when no active antibiotics were prescribed^5^.

However, current culture-based methods used to detect and identify agents of infection are inadequate and slow. Incubation times of 24-48h are necessary to capture the most culturable bacteria associated with the disease. Additional time is required for pathogen ID (2-4h) and in the case of expected AMR for antibiotic susceptibility testing (AST) (18-24h), although for blood cultures, this time has been recently reduced to 4-6 hours using RAST method developed by EUCAST^6^. Thus, the time interval from collecting the patient sample at the ward until the information is available on the antibiotic susceptibility profile is in the best case 2-3 days in the clinical routine.

A plethora of molecular techniques have been developed to reduce the time needed to identify infectious agents and their resistance profiles, including PCR-based or microarray-based technologies^7^. Although they save around 24-48 hours compared to classical AST, they only target known/expected organisms and resistance mechanisms and thus a panel of tests should be performed on each isolate risking that less common pathogens or untargeted resistance mechanisms remain unnoticed. Whole genome sequencing (WGS) can overcome this problem since there is no need for targeted primers/probes to be used. With the rise of real-time sequencing and its affordability, WGS becomes a potent alternative to time-consuming culture-dependent traditional methods. We have recently demonstrated that using Oxford Nanopore Technologies (ONT) MinION and Flongle sequencing platform, infectious agent and its resistance profile can be identified within 10 min – 1 hour after the start of sequencing^8^. However, around 3 hours is still required to prepare the sample for sequencing.

Raman spectroscopy can provide a rapid alternative and overcome many of the problems associated with the current techniques. Since it is culture-free, Raman spectroscopy allows results to be obtained in minutes, rather than several hours and can thus be significantly faster. Unlike ELISA, mass spectrometry, infrared spectroscopy and fluorescence-based techniques, Raman spectroscopy is reagentless and avoids complex sample-preparation steps. Like infrared spectroscopy, Raman uses light to probe the molecular vibrations within the sample, generating a specific molecular ‘fingerprint’ that can be used to identify molecular, biotic, and abiotic components within a sample. However, infrared spectroscopy is highly sensitive to the presence of water, which is ubiquitous in biological samples. In contrast, Raman is highly water-insensitive, offering it an advantage over its sister technique.

Previously, Raman spectroscopy has been used to examine a wide range of biological samples, including an array of microbiological samples in clinical settings. Kloß *et al* used Raman spectroscopy to characterize several respiratory pathogens at the species level^9^, and Ghebremedhin *et al* also used the technique to achieve differentiation of 31 clinical isolates of *A. baumanii* at the strain level^10^.

Surface-Enhanced Raman Spectroscopy (SERS) is a variation of Raman spectroscopy that offers advantages over spontaneous Raman spectroscopy in terms of speed and an improved limit of detection. In SERS, Raman signals are enhanced by electromagnetic and/or chemical interactions with nanoscale metallic structures, such as gold or silver nanoparticles. SERS has been used to distinguish isolates of *E. coli* based on their sensitivity to carbapenem antibiotics^11^, and to identify pathogens common in Cystic Fibrosis sufferers in pellets with silver nanoparticles^12^. SERS has also been used to map colonies of *Pseudomonas aeruginosa* via laser scanning and tracking of a *P. aeruginosa* biomarker^13–17^ Despite being sensitive and fast, SERS has several drawbacks. Most importantly it requires the introduction of exogenous nanomaterials for signal enhancement. The nanostructures must have consistent enhancements, and interact extremely closely and reproducibly with the analyte, which can be difficult to control. Resonance Raman Spectroscopy (RRS) overcomes the requirement for exogeneous materials to provide signal enhancement. RRS utilizes the principle that the cross-section of Raman-active modes varies with wavelength, and improves significantly, as the excitation nears pre-resonance or resonance with an electronic state of the sample. RRS has been used to detect cytochrome *cd1* in bacteria^18^, and UV resonance has been utilized to study both endospore biomarkers and whole bacteria^19^. Grun *et al* studied the possibility of using multiple excitation wavelengths to generate 2D spectra^20^. Their method relied on the use of a pulsed Ti:sapphire laser to generate a tunable excitation from 700-940 nm. This light was converted to third or fourth harmonics to yield excitation from 210-280 nm. Spectra were recorded at 30 wavelengths per species, with approximately 1 minute of switching time between excitations.

Recently we reported the method of multi-excitation Raman spectroscopy (ME-RS) for the strain-level detection of pathogens^21^. This method combined the use of multiple Raman spectra obtained with different wavelengths to interrogate bacterial samples. The natural variations in peak intensity ratios due to the dependence of the Raman cross-section on wavelength gives a more information-rich dataset with only a few minutes of additional analysis time. The combined information was processed by multivariate analysis, in the form of a support vector machine (SVM). This ME-RS approach performed better compared to use of single-excitations individually, and offered highly accurate, strain-level classification, even inside a complex artificial sputum media (99.75% accuracy). Further, an accuracy of 100% was obtained for differentiation of methicillin-resistant and methicillin-sensitive *S. aureus* strains.

Here, we apply our newly developed multi-excitation Raman spectroscopy^21^ method to the identification of clinical isolates of four WHO priority pathogens (*E. coli, K. pneumoniae, S. aureus* and *A. baumannii*) with known antibiotic susceptibility phenotypes and resistomes^8,22–24^. Spectra were concatenated and used to train a Support Vector Machine (SVM), resulting in highly accurate strain and species identification of the isolates. This work demonstrates the potential for ME-RS technique as a rapid first-stage method for informing the prescription of appropriate antibiotics prior to more extensive confirmatory lab-based testing by WGS. A combined workflow with ME-RS and WGS could be transformative for the detection and identification of infections and AMR in human and animal health.

## Methods

### Dataset description

Wild-type (-WT) and phenotypically resistant (-R) strains of Gram-negative *Acinetobacter baumannii* (INN-WT, K55-33-R), *Escherichia coli* (CCUG17620-WT; A2-39-R), *Klebsiella pneumoniae* (225-R) and Gram-positive *Staphylococcus aureus* (NCTC8325-WT; CCUG35600-R) were used for the study. Phenotypic information on *E. coli, K. pneumoniae* and *S. aureus* strains was previously published in^8,22–24^. Minimum inhibitory concentration (MIC) values of *Acinetobacter* strains against ciprofloxacin, gentamicin, meropenem and colistin were determined by broth microdilution using Sensititre surveillance EUVSEC 96 well plates (ThermoFisher, USA) as described in^23^. Isolates were classified as susceptible (wild type) and resistant according to the European Committee on Antimicrobial Susceptibility Testing (EUCAST) Breakpoints v 12.0 (December 2021).

### Genome sequencing & Bioinformatics for AMR and resistance

Genomic background of *E. coli*, *K. pneumoniae* and *S. aureus* used for this study was extensively characterized in our previously published work^8,22–24^. In brief, isolates were sequenced using either Illumina^23^, ONT (MinION and Flongle)^8,22^ or both^24^. Regardless of the technology used, sequencing data were filtered for quality and length, and subsequently assembled using either SPAdes v 3.13.1 (Illumina) or Unicycler v 0.4.9 (ONT MinION). In case data from both technologies were available, hybrid genome assembly was performed using Unicycler v 0.4.9.

*A. baumannii* isolates were processed as described in^23^, sequenced on a MiSeq Illumina platform using MiSeq v3 chemistry. Sequencing reads were demultiplexed, quality-filtered and assembled using SPAdes v 3.13.1 following previously published protocol^23^.

Genome assemblies were searched for ARGs using Abricate v 1.0.1 which performs mass screening against multiple ARG databases (NCBI, ARG-ANNOT, ResFinder, MEGARES). Eighty percent identity and 80 % query coverage were used as cutoffs. Plasmids were searched against PLSDB database using 90 % identity as a cutoff^25^.

### Bacterial culture

Isolates were kept in 25 % glycerol stock at −80 °C. A loop-full of each isolate stock was streaked on BHI agar plates and cultured overnight at 37 °C. After visual confirmation of colonies’ homogeneity (shape, size, color), 5-10 colonies were picked and transferred to 1.5 ml BHI broth and cultured overnight at 37 °C. After confirmed growth (OD600 >1.0), 500 μl of bacterial culture were transferred to 500 μl of 50 % glycerol solution. The stock was then sent to the Raman spectroscopy lab on dry ice, where they were kept at −20 °C until the experiment.

BHI medium was prepared from the BHI powder (VWR, USA), following the manufacturer’s protocol. 15 g/l of agar was added to the medium for preparing the plates.

### Raman microspectroscopy

To prepare samples for spectroscopic analysis, bacterial cultures were washed three times in deionized water by centrifugation (4000g, 10 minutes) in a Heraeus Megafuge centrifuge. The resulting pellet was applied to a fused quartz slide (UQG Optics, UK), and dried with gentle heating.

Raman microspectroscopy experiments were conducted using a Renishaw InVia Raman microscope (Renishaw, UK), with a Leica DM 2500-M bright field microscope and an automated 100 nm-encoded XYZ stage. The samples were excited using 532 nm and 785 nm lasers directed through a Nikon 100× air objective (NA = 0.85), with collection after a Rayleigh edge filter appropriate to each excitation wavelength, and a diffraction grating (532nm: 1600 L/mm, 785nm: 1200 L/mm) that dispersed the Raman-scattered light onto a Peltier-cooled CCD (1024 pixels × 256 pixels). Calibration of the Raman shift was carried out using an internal silicon wafer using the peak at 521 cm^−1^. Spectra were acquired over three accumulations of 5 s each.

### Spectral data processing and chemometric analysis

All spectra were cleared of cosmic rays prior to analysis using Renishaw Wire 5.1 software and then imported into iRootLab version 0.17.8.22-d for Matlab28 for further processing. Spectra were truncated to the 600-1600 cm^−1^ spectral region and then background subtracted, wavelet denoised to smooth them, and normalized to their maximum intensity. Concatenation of multi-excitation spectra was performed by appending the 532nm spectrum to the end of the 785nm spectrum. Wavenumber variables were changed to integer ‘observation’ values to prevent issues arising from having multiple intensity values at each wavenumber.

Two-hundred and ten spectra were used to train the SVM classified (30 for each strain), and the same number of spectra was used for validation. For this study, iRootLab’s in-built Principal Component Analysis (PCA), SVM, and k-fold cross-validation functionality was applied to the processed spectra to classify the bacteria by strain. In SVM, the default iRootLab parameters for c and gamma (c = 1, gamma = 1) were used. The analytical process is schematically summarized in Figure 1. The full step-by-step methods for the SVM have been included as Figure S1.

**Figure 1:**
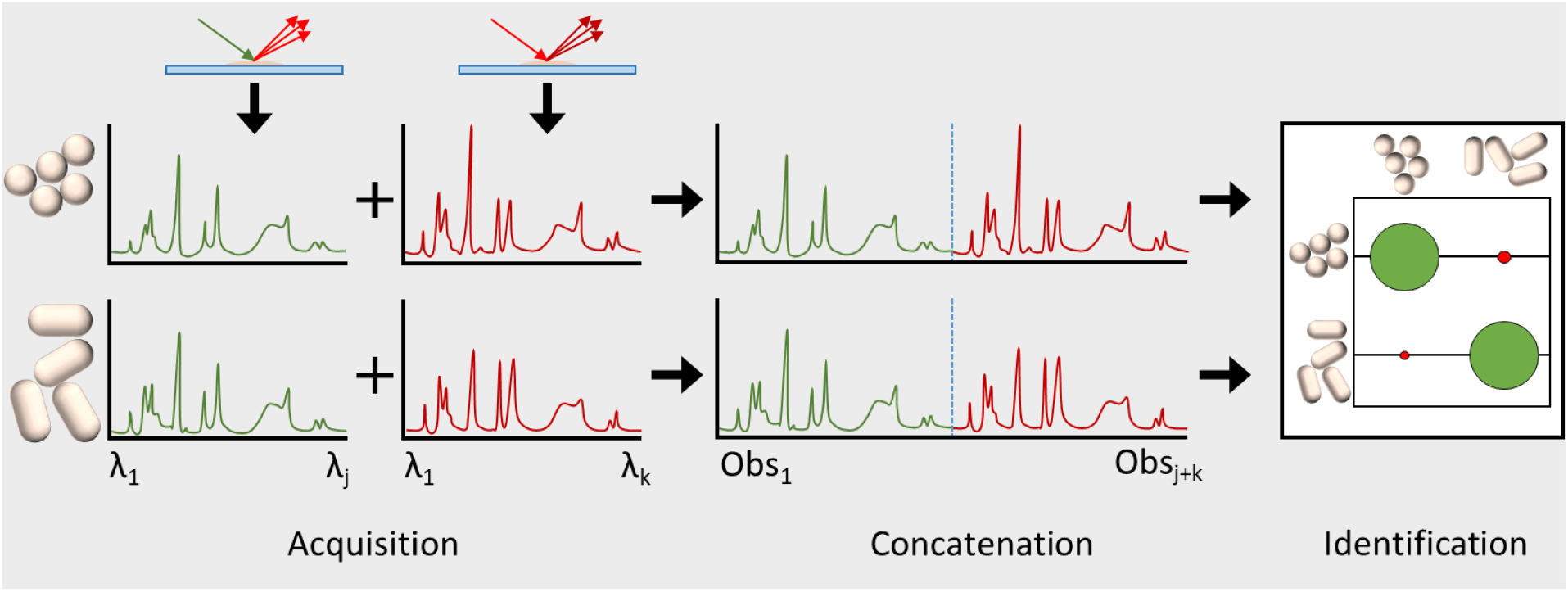
Diagrammatic representation of the workflow for spectral analysis. Spontaneous Raman spectra are recorded at two wavelengths, and then concatenated onto one another. Raman shift (wavelength or wavenumber) variables are replaced with a positive integer, termed an observation. The combined data of intensity vs observational variables is then fed into a support vector machine for training and testing purposes.

## Results and Discussion

### Sequencing

Wild type strains of *E. coli* (CCUG17620) and *A. baumannii* (INN) did not possess any antimicrobial resistance genes, whereas the *S. aureus* NCTC 8325 genome contained genome encoded *fosB* gene. Resistant strains, on the other hand, possessed plasmids with a wide variety of genes conferring resistance towards penicillins, 3^rd^ generation cephalosporins, carbapenems, tetracyclines, fosfomycin and aminoglycosides (Table 1). In methicillin-resistant *S. aureus* CCUG35600, ARGs were chromosomally encoded (Table 1).

**Table 1:**
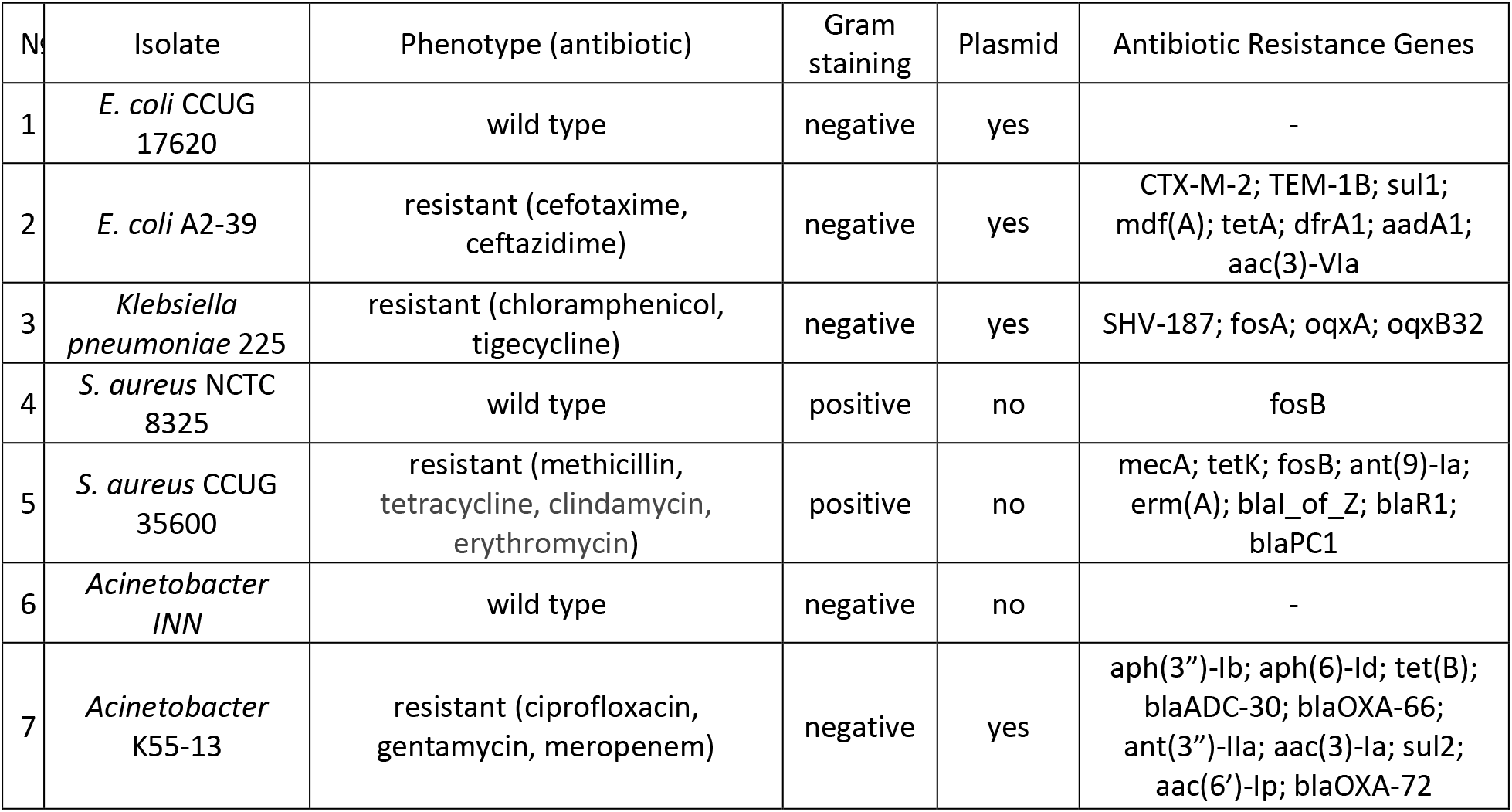
Overview of isolates’ phenotype and genomic background.

Twenty percent of *K. pneumoniae* 225 genome aligned to *E. coli* CCUG17620 and an average nucleotide identity (ANI) between these isolates was 84.6%. In case of *E. coli* A2-39, 18.6 % of *K. pneumoniae* 225 genomes were aligned and ANI was found to be 84.4 %.

### Raman spectral data

Using our previously reported method for multi-excitation Raman spectroscopy^21^ or ME-RS, spectra of seven strains of WHO priority pathogens were recorded at 532 nm and 785 nm excitation. Class mean spectra (*n* = 30) for both excitations are presented in Figure 2. In both cases, there is a large degree of spectral similarity between the different classes, except for *S. aureus 8325* at 532 nm excitation. This strain exhibits large peaks at 1157 cm^−1^ and 1525 cm^−1^, which are associated with the conjugated - C=C-backbone of carotenoids^26,27^. These molecules are pre-resonantly excited at wavelengths around this region^28,29^. Several other peaks can be observed across both spectra. Within the 532 nm spectra, we assign the 747 cm^−1^ peak to the presence of DNA^30^, along with the 781 cm^−1^ peak, which arises from ring breathing modes of cytosine^31^. The peak at 1004 cm^−1^ arises from the ring breathing mode of phenylalanine^32^. The sharp peak at 1128 cm^−1^ contains contributions from lipid C-C modes and C-N stretches in protein^33,34^. Lastly, we attribute the feature at 1585 cm^−1^ to olefinic C=C modes in proteins^35^. Within the 785 nm spectra, many of the features of the 532 nm spectra are preserved, but additional peaks are also seen. We ascribe the feature at 1030 cm^−1^ to a mixture of C-C and C-H modes in phenylalanine^32,35^. Lastly, we attribute the broad peak around 1450 cm^−1^ to a complex mixture of C-H modes associated with molecules such as proteins, lipids, and nucleic acids^34,36–38^.

**Figure 2:**
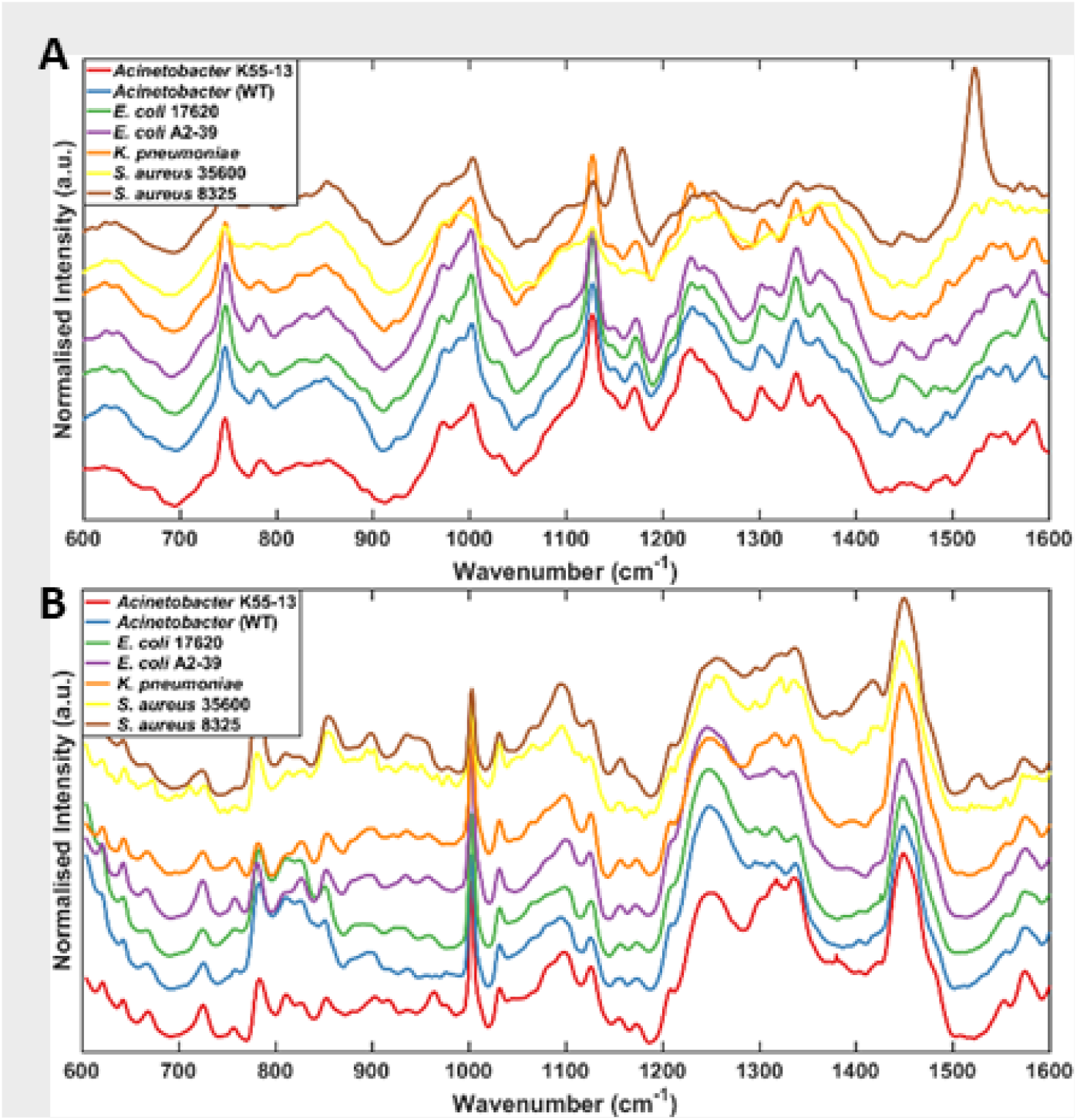
Normalized class means (offset for clarity) of the Raman spectra for the 7 bacterial strains used in this study, taken using **(A)** 532 nm excitation, and **(B)** 785 nm excitation. (*n* = 30) The presented spectra illustrate the changes in peak intensity ratios that arise from changes in the excitation wavelength.

To determine the ability of the multi-excitation method to elucidate several biologically relevant characteristics of the WHO pathogens and to also see their correlation with the sequencing information, we utilized SVM with ten-fold cross validation, and performed PCA as a point of comparison. Initially, we analysed the samples based on their species and the presence or absence of a thick peptidoglycan layer in the cell wall (i.e., Gram-positive versus Gram-negative). The results for the SVM analyses are shown in Figure 3 A&B. Using this method, we achieved 100% accuracy for the differentiation of Gram-positive and Gram-negative bacteria. PCA for Gram-positive and Gram-negative bacteria also showed clear separation of the two classes along PC1 (Figure S2 B), with no overlap of the 95% confidence intervals of the groups. It is a very promising result since *Acinetobacter* is known to be occasionally falsely classified as a Gram-positive bacterium^39^, which can impair proper infection treatment^40^.

**Figure 3:**
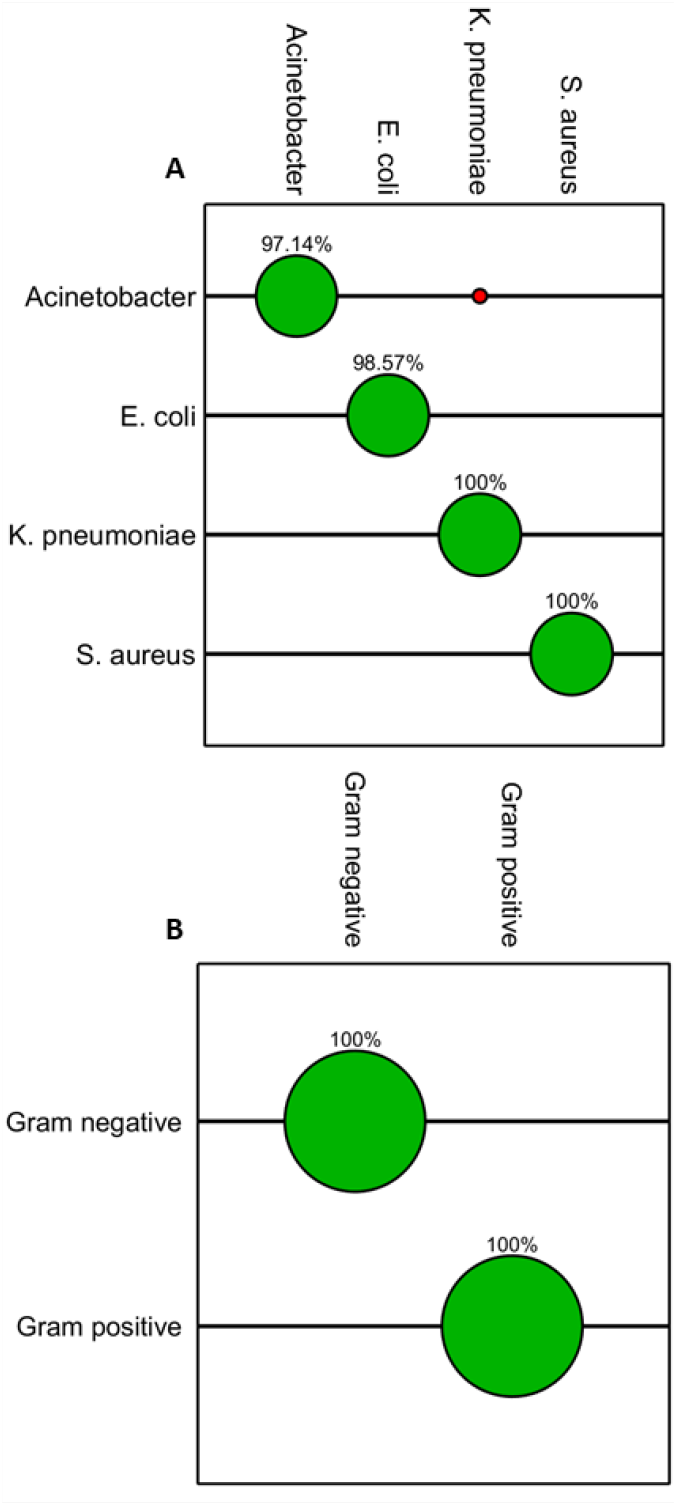
**(A)** Classification accuracies for SVM species-level identification of the bacterial strains used in this study. **(B)** Classification accuracies for SVM delineation of gram-positive and gram-negative bacteria. The size of green circles corresponds to the number of correct identifications for that species. Red circles indicate incorrect classifications. A total of 210 concatenated ME-RS spectra were input into SVM.

A mean classification accuracy of 99.93% for species-level classification was also achieved by the SVM analysis of ME-RS data. 2 out of 60 *A. baumanii* spectra were incorrectly classified as *K. pneumoniae*, and 1 of 60 *E. coli* spectra were identified as *A. baumanii*. In comparison, the species-level PCA exhibited significant overlap of the 95% confidence intervals of all species other than *S. aureus* (Figure S2 A). *S. aureus’* separation from the other species is congruent with the species’ structural differences in its cell wall. Similarly, all inter-species classification errors in the SVM analyses occurred between Gram-negative species.

We also attempted to differentiate the bacteria according to their drug resistance phenotype (i.e., resistant versus wild type), and to classify each strain in the study as a unique class. For this proof-of-concept study, we only separated bacteria into resistant (plasmid-mediated or chromosomal resistance to any antibiotic) and wild type (susceptible to all antibiotics or chromosomal mediated resistance). A mean accuracy of 99.45% was achieved across the two classes, with all misclassifications occurring in the wild type class (Figure 4). Whilst any misclassification is undesirable, it is preferable that any misclassification be a false positive for drug resistance rather than a false negative, which may have negative consequences in terms of adverse outcomes for patients or animals. On the other hand the PCA plot of the phenotype information revealed extensive overlap of the wild type and resistant classes (Figure S3 A). Interestingly, although *S. aureus* NCTC 8325 has fosfomycin resistance gene, it was still separated from *S. aureus* CCUG35600 MRSA strain. The same was true for *E. coli* isolates, where spectra from an ESBL-positive strain were separated from a wild-type *E. coli* strain. Unfortunately, wild type *K. pneumoniae* isolate was not included in this study. *K. pneumoniae* are intrinsically resistant to ampicillins, and it will be important to include a wild-type *K. pneumoniae* strain in the follow-up experiments. At this stage, our experimental data is rather limited, and it is yet to be established whether ME-RS can distinguish between resistance to a given antibiotic. Nevertheless, this study clearly highlights that ME-RS has the potential to be used as a rapid antibiotic susceptibility assessment tool.

**Figure 4:**
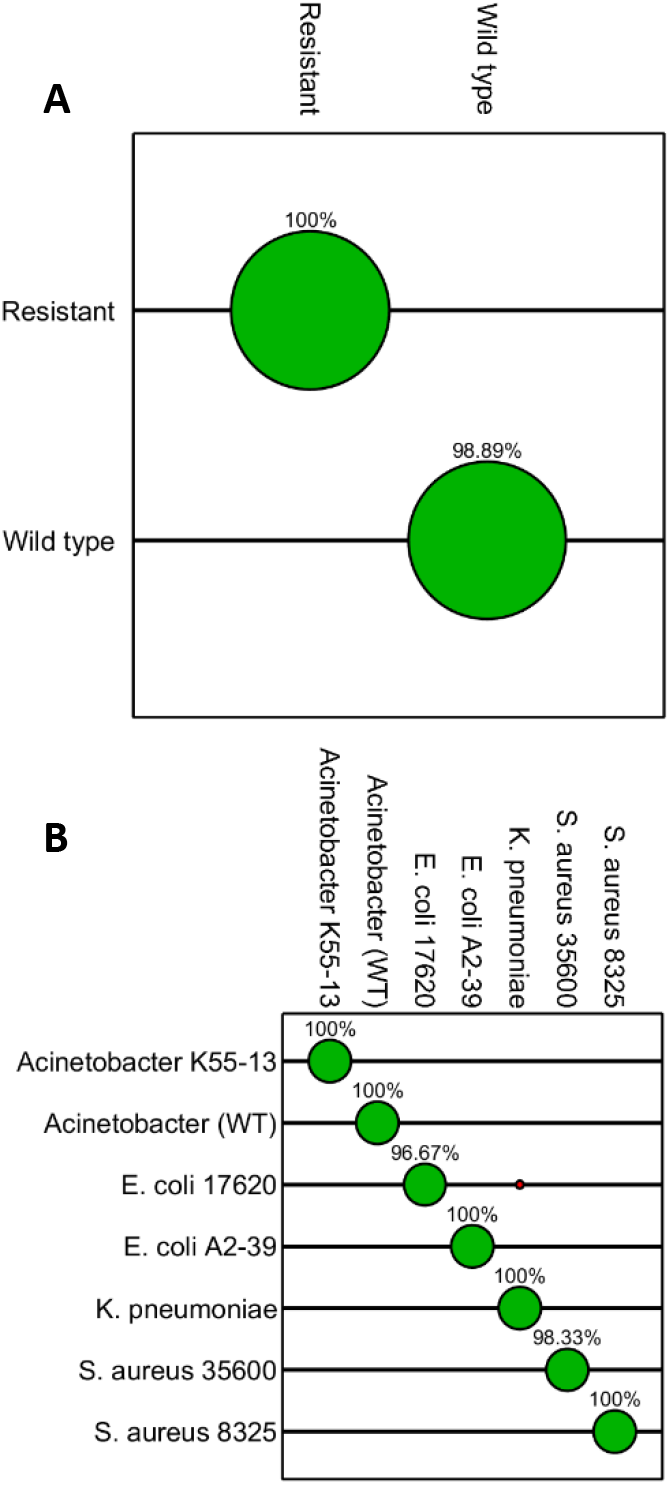
**(A)** Classification accuracies for SVM delineation of bacteria exhibiting drug sensitivity and those exhibiting drug resistance. **(B)** Classification accuracies for SVM strain-level identification of the bacteria used in this study. The size of green balls corresponds to the number of correct identifications for that species. Red balls indicate incorrect classifications. A total of 210 spectra with 30 spectra of each strain were input into SVM.

Lastly, we tested our method as a means of classifying each strain separately and performed a PCA for comparison. The results of this analysis are shown in Figure 4 B and Figure S3 B. In the PCA space, *S. aureus* is well resolved from the Gram-negative species used in the study, with both strains also being resolved from one another, suggesting that even PCA can differentiate drug-resistant and drug sensitive strains in this species; however, the Gram-negative species exhibit sizeable overlap between both species and strains. Clearly, this is highly undesirable. This strain-level classification essentially represents a simultaneous classification along both phenotype and species lines, and the SVM model performed extremely well, achieving a mean classification accuracy of 99.29%. Within this model, *A. baumannii* strains were classified correctly 100% of the time, as were *S. aureus* NCTC8325 and *E. coli* A2-39. 3.33% of *E. coli* CCUG17620 was misclassified as *K. pneumoniae* 225, and 1.67% of *S. aureus* CCUG35600 was misclassified as *S. aureus* NCTC8325. *E. coli* CCUG17620 had 20.3 % of its genome aligned to *K. pneumoniae* 225 genome, whereas *E. coli* A2-39 shared 18.6 %. Average nucleotide identity (ANI) of these strains to *K. pneumoniae* equaled 84.6 % and 84.4 % respectively. Both the *E. coli* and *S. aureus* misclassifications result in a misidentification of the sample’s drug-resistance phenotype, which is undesirable; however, as both sets of misclassifications incorrectly identified the sample as drug-resistant, the clinical decision to change to a different antibiotic to circumvent resistance is unlikely to be problematic in treating an infection. Interestingly, comparison of the species and strain level classifiers suggests that underlying phenotype may influence species identification. In the species-level classification, *A. baumannii* was misidentified as *K. pneumoniae* in a small number of instances. This misclassification, along with misclassifications of *E. coli* as *A. baumannii* disappeared when classifying strains independently. It is possible that this is caused by physical changes arising from genetic differences between the strains, which may confound the more general species classifier. Given this, and the importance of AMR in the modern clinical setting, it may be preferable to classify unknown samples along strain lines.

Previously, Raman spectroscopy has been used with single excitation wavelengths to detect pathogens from patient samples, but exposure times involved in the analysis required photobleaching steps of 15-30 minutes to allow Raman spectral features to become prominent enough to achieve an accuracy above 95%^41^. As we have previously reported, our methodology provides clear spectra in around a minute for classification of two species, even in artificial sputum^21^. Here, we have extended the previous work to more species of clinical significance with similar levels of accuracy. Ho *et al* utilized deep learning to identify pathogens based on their Raman spectra, and reported an accuracy if 99.7%, which is comparable to the accuracies we obtain in this work.

## Conclusions

Here, we have demonstrated the extension of our multi-excitation Raman spectroscopy method to four WHO priority pathogens with known antibiotic susceptibility phenotypes and resistomes. Multi-excitation Raman combined with multivariate analysis was applied to a variety of biologically relevant classification problems. Mean classification accuracies for species, Gram staining, drug resistance phenotype, and strain were 99.93%, 100%, 99.45%, and 99.29%, respectively, which is consistent with previously reported findings. The classification by ME-RS is well supported by the detailed phenotypic and gene resistance profiling data. We observe a small (<1%) number of misclassifications which are explained by phenotypic differences and genome alignment. These findings demonstrate the utility of our method to assist in the identification of a range of WHO priority pathogens, and to provide relevant information about microbiological samples, which can later be verified by genomics or conventional microbiological assays. Our results establish the potential of ME-RS as a rapid first-stage analytical tool that can complement WGS for phenotype prediction and resistome analysis. Such a workflow can be hugely impactful to handle and prevent the spread of AMR and could lead to potential future use in clinical microbiology.

## Acknowledgements

This research was funded by the Norwegian research council, grant number 273609, to AMR-Diag, and to the internal strategic research grant 2021 from the Inland Norway University of Applied Sciences. The authors would like to thank Anne Bergljot and Arne Taxt for help with the selection of the bacterial isolates and Stephan A. Frye for performing the WGS. SM, AL, GD and NH acknowledge funding from EPSRC grant EP/T020997/1 and EPSRC Impact Acceleration Account (IAA) award. JSW and CJH acknowledge funding from the BBSRC and Innovate UK, IKC National Biofilms Innovation Centre BB/R012415/1.

