## Supplementary Information and Figures for "Multi-excitation Raman Spectroscopy Complements Whole Genome Sequencing for Rapid Detection of Bacterial Infection and Resistance in WHO Priority Pathogens"

Figure S1: Workflows for PCA, SVM, and PCA-SVM in iRootLab

**Step-by-step multivariate statistical methods performed in iRootLab (v0.17.8.22-d)**

### Support Vector Machine

1. Load combined dataset

2. (If desired), select 'Pre' and polynomial baseline correction (5$^{th}$ order polynomial, epsilon = 0, no background spectra, index of background [1])

3. Select 'Pre' and wavelet de-noising. Use default settings (wavelet name: 'haar', 6 decompression levels, thresholds: [0, 0, 0, 100, 1000, 1000])

4. Select 'Pre' and normalization. Use default settings (Normalized to maximum intensity)

5. Select 'Sub-dataset Generation Specs' on the left
6. Create a new K-fold cross-validation (used 10-fold)
7. Select 'Block' on the left
8. Create a new support vector machine, using the default parameters (c=1, gamma =1. iRootLab will alter these during the analysis).

9. Return to 'Dataset' on the left and select your pre-processed data.

10. Select 'AS' and Grid Search.

11. In the menu that pops up, use the following parameters: a. SGS - SGS_crossval01 b. Classifier – clssr_svm01 c. Templates: SVM d. Test post-processor, chooser, and estimation post-processor are left as default settings.

12. Allow the grid search to run, then select 'Log' on the left

13. Select log_gridsearch_gridsearch01 and extract_block S3

14. Return to 'Dataset' and import your validation data

15. Select the validation data and select 'AS' and then Rater.

16. In the menu that pops up, use the following parameters: a. Classifier - Clssr_svm01 b. SGS - SGS_crossval01 c. As before, leave other parameters as the defaults

17. Allow to run, then select 'Log' and estlog_classxclass_rater01.

18. Select confusion matrices and hit okay to retrieve the results.

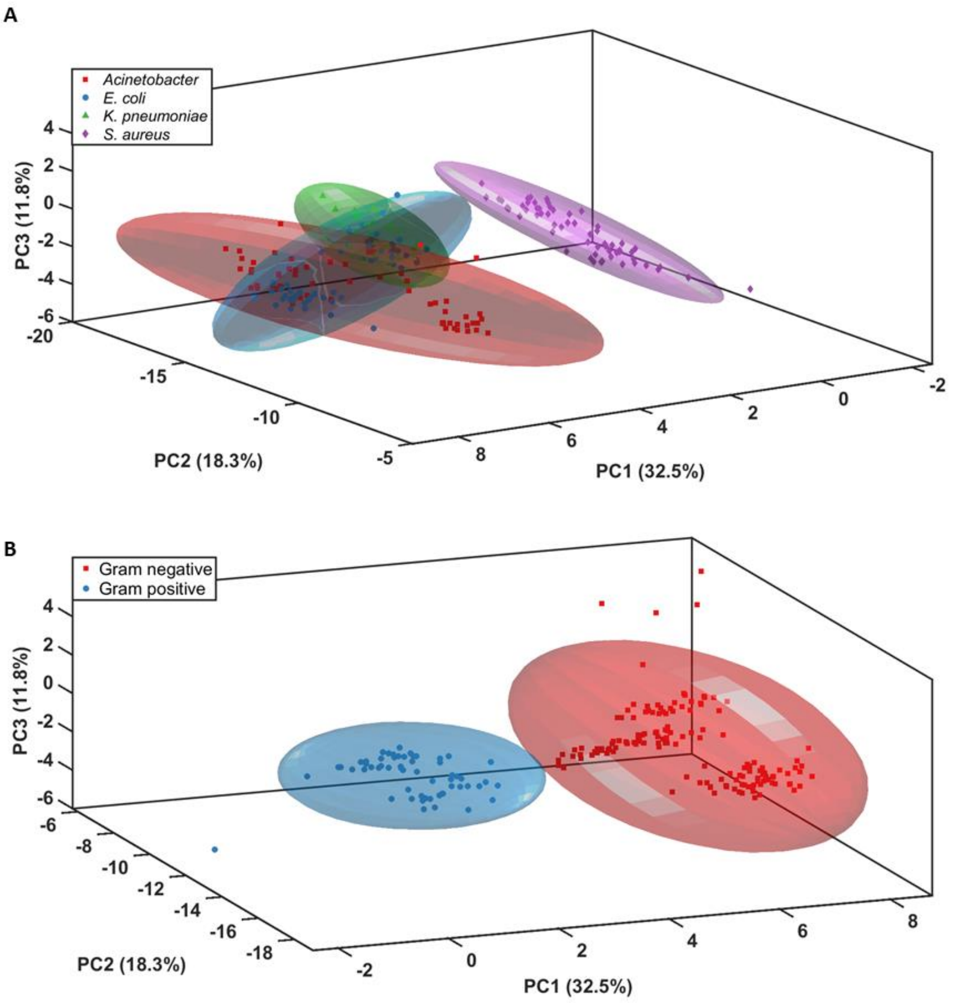


**Figure S2:** Projections of the first three principal components for the spontaneous multi-excitation Raman spectra of the four bacterial species used in this study, showing **(A)** Separation at the species level and **(B) s**eparation of gram-positive and gram-negative bacteria. Colored ellipses represent the 95% confidence intervals in both cases.

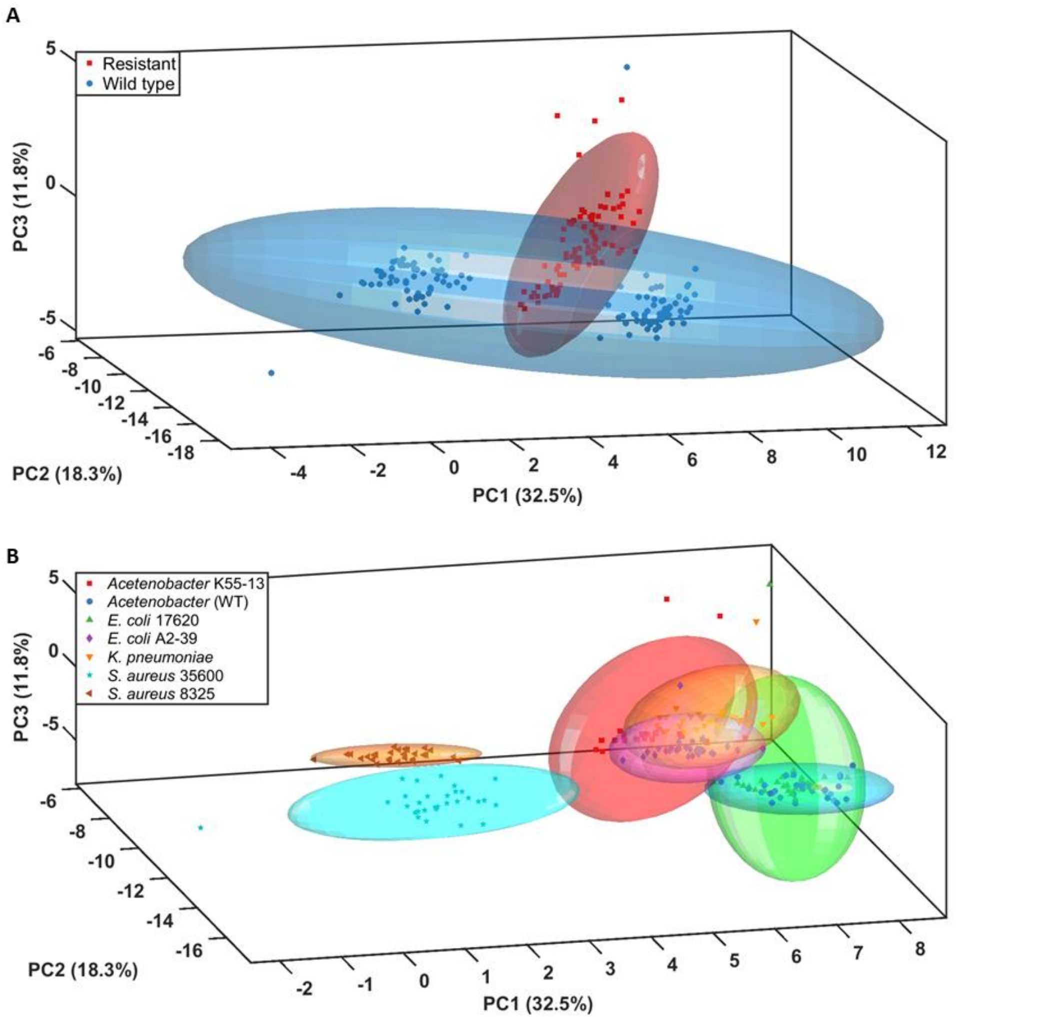


**Figure S3:** Projections of the first three principal components for the spontaneous multi-excitation Raman spectra of the four bacterial species used in this study, showing **(A)** Separation of drug resistant and drug sensitive strains, and **(B)** separation of the four species of bacteria at the strain level. Colored ellipses represent the 95% confidence interval in both cases.
